# *Ab initio* RNA structure prediction with composite language model and denoised end-to-end learning

**DOI:** 10.1101/2025.03.05.641632

**Authors:** Yang Li, Chenjie Feng, Xi Zhang, Yang Zhang

## Abstract

RNA structures are essential for understanding their biological functions and developing RNA-targeted therapeutics. However, accurate RNA structure prediction from sequence remains a crucial challenge. We introduce DRfold2, a deep learning framework that integrates a novel pre-trained RNA Composite Language Model (RCLM) with a denoising structure module for end-to-end RNA structure prediction. DRfold2 achieves superior performance in both global topology and secondary structure predictions over other state-of-the-art approaches across multiple benchmark tests. Detailed analyses reveal that the improvements primarily stem from the RCLM’s ability to capture co-evolutionary pattern and the effective denoising process, leading to a more than 100% increase in contact prediction precision compared to existing methods. Furthermore, DRfold2 demonstrates high complementarity with AlphaFold3, achieving statistically significant accuracy gains when integrated into our optimization framework. By uniquely combining composite language modeling, denoise-based end-to-end learning, and deep learning-guided post-optimization, DRfold2 establishes a distinct direction for advancing *ab initio* RNA structure prediction.

## Introduction

Understanding the structure and function of RNA molecules has been a central focus of molecular biology and the pharmaceutical industry. RNAs, particularly non-coding RNAs, fold into specific structures that can be functional in various cellular processes, including gene regulation (e.g., transcription and translation), catalysis, cellular signaling, stress responses, and genome defense ^1^. Due to their functional significance, RNAs have emerged as promising targets for small-molecule drug development, especially for human diseases that lack traditional protein targets ^2, 3^. With the exponential growth of RNA sequence data from high-throughput sequencing and the widening gap between known sequences and experimentally resolved RNA structures, determination of atomistic structures of RNAs from the primary sequences has become increasingly urgent.

Researchers have developed multiple approaches to study RNA structures, ranging from chemical probing ^4^ and proximity ligation methods ^5^ for detecting base pairs or other interactions, to structural biology techniques such as X-ray crystallography ^6^, NMR spectroscopy ^7^, and cryo-EM ^8^. Despite higher resolution, experimental RNA 3D structure determination is often prohibitively expensive and sometime infeasible. Consequently, computational approaches have faced growing demands for high-quality 3D structure modeling directly from sequences.

Computer-based RNA structure prediction has evolved considerably over the past decades. Traditional template- and fragment-based approaches ^9–12^ construct structure models using known templates or fragments but are constrained by the limited availability of experimentally solved structures. *Ab initio* methods ^13–15^ attempt to predict structures from sequence alone; however, traditional physical or statistical force fields often lack the accuracy and specificity needed to fold RNAs with complex topologies. Recent advances in deep learning have introduced several innovative approaches. For example, DeepFoldRNA ^16^ and trRosettaRNA ^17^ utilize transformer based models to predict local geometries and reconstruct coordinates from the predicted restraints. On the other hand, end-to-end models like RhoFold ^18^ and RoseTTAFoldNA ^19^ directly predict 3D coordinates, inspired by AlphaFold2 ^20^ in protein structure predictions. Our previous approach, DRfold ^21^, combines both end-to-end and geometry-based predictions through optimizing a hybrid potential function with a differentiable fold program ^22^.

Despite the progress, RNA structure prediction accuracy has not yet reached a so-called “AlphaFold moment” ^23^, partly due to limited solved structures compared to proteins. Language models have shown promise in protein structure prediction by capturing key features like co-evolution patterns and improving sequence representation from massive unsupervised sequence data ^24^. Their ability to predict structures from single sequences is particularly valuable for RNAs, while multiple sequence alignment (MSA) construction ^25^ is more resource intensive and the usefulness often varies depending on the depth of MSAs.

In this study, we introduce DRfold2, a new deep learning-based framework for high-quality RNA structure prediction. The core novelty of DRfold2 consists of a pretrained RNA Composite Language Model (RCLM), which improves likelihood approximation and captures co-evolutionary signals from unsupervised RNA sequences more effectively than the previously used embeddings learned from structure prediction pipeline ^21^. Next, the sequential and pairwise representations from RCLM are processed by an RNA Transformer-based pipeline, powered with a denoising module for end-to-end structure and geometry prediction. A post-processing protocol is then implemented for final RNA model selection and optimization, built on the end-to-end and geometry potentials. The large-scale benchmark results demonstrate that co-evolution features learned by RCLM from unsupervised data, when integrated with denoising structure learning, significantly enhance the accuracy of *ab initio* RNA structure prediction.

## Results

The overall DRfold2 pipeline for RNA structure prediction is illustrated in **Fig. 1A**. Starting from an RNA sequence, DRfold2 first employs a pretrained RCLM to embed the query into sequence and pair representations. Here, RCLM achieves superior sequence pattern recognition through training on large-scale unsupervised sequence data using a composite likelihood^26^ maximization approach (**Fig. 1B**). These sequence and pair representations are processed by a set of RNA Transformer Blocks (**Fig. 1C**), which produce necessary representations for structure folding. DRfold2 then generates RNA conformations through a Denoising RNA Structure Module (DRSM) in an end-to-end fashion (**Fig. 1D**). The final RNA models are obtained through a post-processing CSOR protocol that is designed to select and refine conformation decoys generated from a pool of checkpoints (**Fig. 1E**).

**Figure 1.**
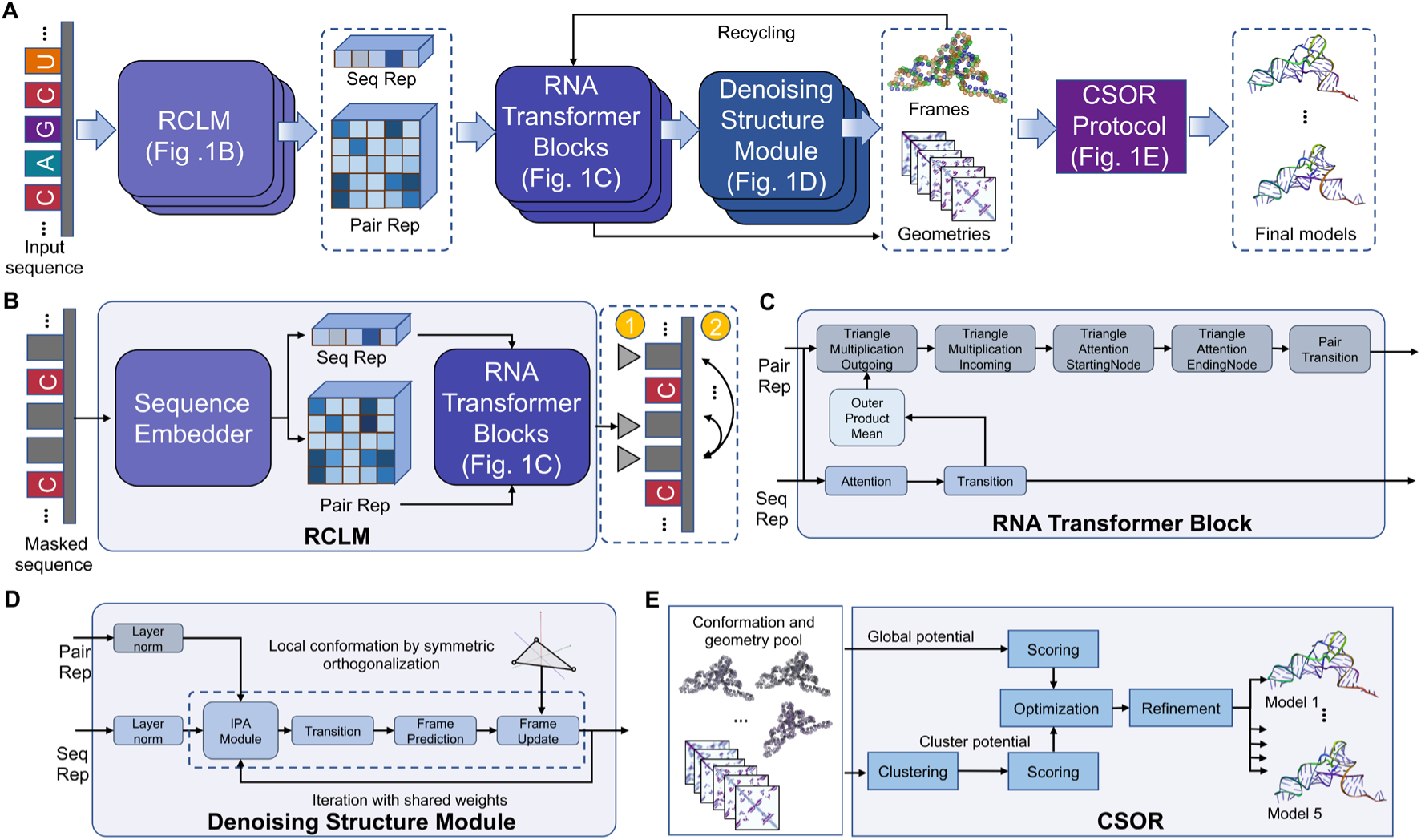
The pipeline of DRfold2 for end-to-end RNA structure prediction. **(A)** Overview of DRfold2 pipeline. **(B)** Details of training RCLM with masked negative composite log likelihood loss function. **(C)** Details of RNA Transformer Block. **(D)** Details of Denoising RNA Structure Module. **(E)** Detailed pipeline of CSOR Protocol as post-process to select and refine final RNA models.

To objectively benchmark the performance of DRfold2, we constructed an independent test dataset containing 28 RNA structures with sequence length < 400 nts collected from (1) recent RNA-Puzzles targets ^27^, (2) CASP15 RNA targets ^28, 29^, and (3) recently released RNA-only structures in Year 2024 (before August 1st) from Protein Data Bank (PDB) ^30^. Notably, large synthetic RNAs from CASP15 were excluded due to their deviation from naturally occurring RNAs, which serve as the primary focus for most functional assays and drug design efforts. To ensure rigorous evaluation, our training set only includes RNA structures released before 2024, while excluding any RNAs sharing more than 80% sequence identity with the test RNAs. Both the training and test datasets are made publicly available at https://zhanglab.comp.nus.edu.sg/DRfold2/data.zip.

### DRfold2 outperforms existing methods in global RNA tertiary structure prediction

We first evaluate the performance of DRfold2 in comparison with five state-of-the-art methods: RNAComposer ^12^, trRosettaRNA ^17^, RhoFold ^18^, RoseTTAFoldNA ^19^, and DeepFoldRNA ^16^. While RNAComposer utilizes a fragment assembly and refinement approach, the other four methods are based on recently developed deep learning models for RNA structure prediction, each with different strategies and focuses. To ensure reproducibility, all methods were installed and executed locally, except for RNAComposer, which we accessed via web server. **Fig. 2A** compares the TM-score and RMSD between DRfold2 and the control methods, under different sequence identity cut-off thresholds from 50% to 80% between the test and training RNAs. Here, the Template Modeling Score (TM-score) ^31, 32^ is a length-independent scoring function to assess the quality of predicted full-length RNA structures, ranging from 0 to 1, with higher values indicating greater similarity to the native. TM-score was designed to address the limitations of the widely used Root Mean Square Deviation (RMSD), which is highly sensitive to local structure derivations. Both TM-score and RMSD are computed on the P-atoms of the predicted and experimental structures.

**Figure 2.**
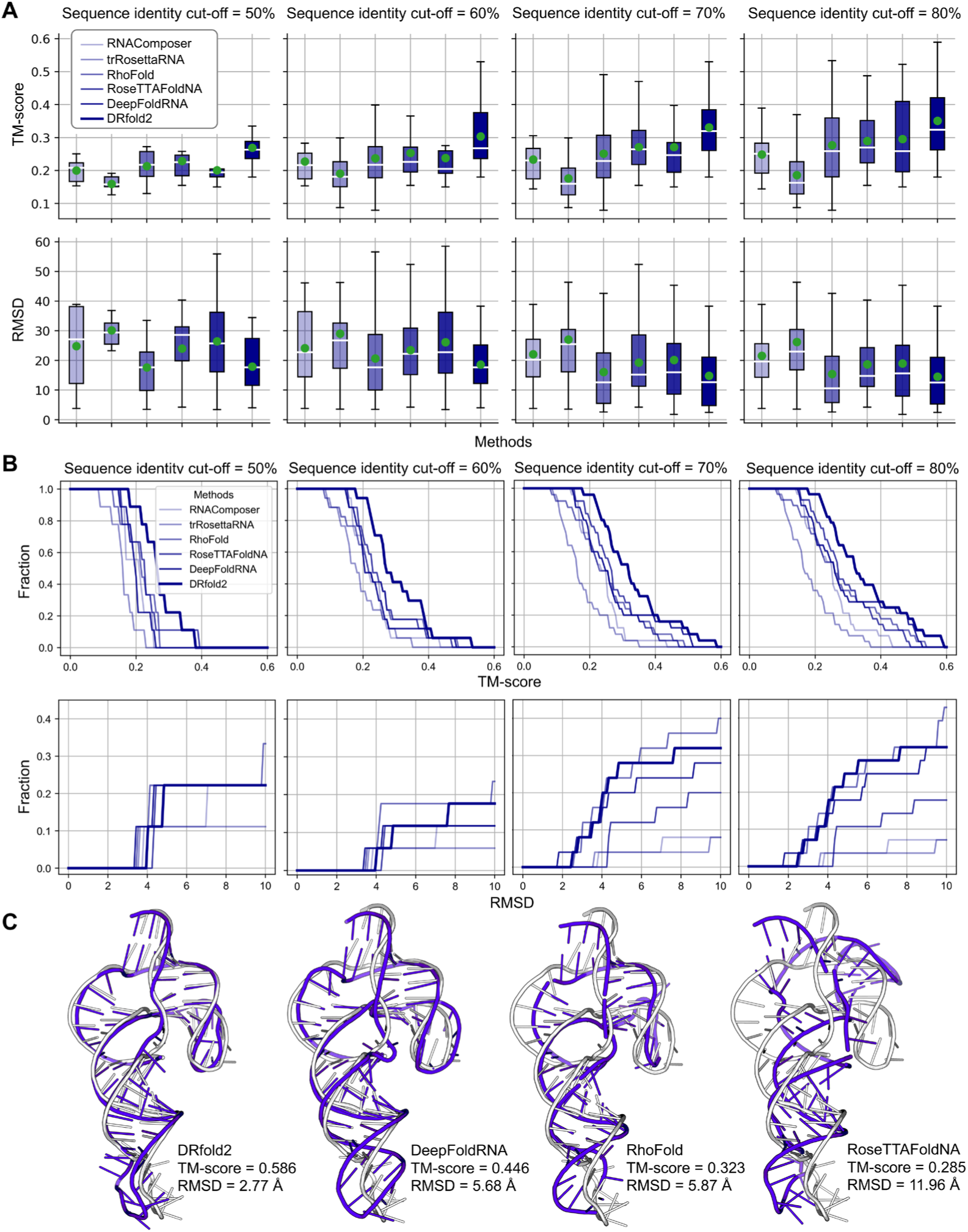
RNA structure prediction results on the test dataset by different methods across different sequence identity thresholds. **(A)** Box plots of TM-score and RMSD for six RNA structure prediction methods evaluated at different sequence identity cutoffs (50-80%) between training and test datasets, with green point and white horizontal line representing the mean and median values respectively. **(B)** Cumulative distribution plots of TM-score and RMSD for the predicted models. Upper panels display the fraction of models above different TM-score thresholds, while lower panels show the fraction of models below different RMSD values. **(C)** A representative modeling example from Chimpanzee CPEB3 HDV-like ribozyme (PDB ID: 7QR3), with models predicted by four better-performing methods (blue cartoons) overlaid on experimental structure (gray cartoons).

Within the test dataset, there are 9, 17, 25, and 28 targets with sequence identities below 50%, 60%, 70%, and 80%, respectively, relative to the training dataset. In general, prediction accuracy of DRfold2 increases for the targets with higher sequence identity to the training set. Notably, this trend holds for all control methods even though no filters was applied to their training dataset, probably because these methods were trained on RNAs released before 2024 which have a similar homologous distance to the new test dataset used in this study. It is observed that DRfold2 consistently achieves the highest average TM-score among all methods across all sequence identity thresholds. For example, at the 80% cutoff, DRfold2 attains an average TM-score of 0.351 which is 18.6% higher than the second-best method, DeepFoldRNA (TM-score=0.296). The difference is statistically significant, with a p-value of 7.18E-04 in the one-sided Student’s t-test, despite the low degree of freedom. Even with the most stringent test subset (50% sequence identity cutoff), DRfold2 achieves an average TM-score of 0.269, which is 17.5% higher than the runner-up, RoseTTAFoldNA (TM-score=0.229). Meanwhile, DRfold2 exhibits a lower RMSD than all other control methods across different sequence identity thresholds.

**Fig. 2B** further shows a comprehensive performance comparison using accumulative fraction plots of TM-score and RMSD. Each curve represents the fraction of predictions achieved better than specific TM-score or RMSD thresholds. DRfold2 (highlighted in dark blue) consistently outperforms other methods in different accuracy ranges and across different sequence identity cutoffs, especially on TM-scores that are less sensitive than local fluctuations. These systematic comparison results suggest that DRfold2 provides robust global tertiary structure predictions across diverse RNA sequences and accuracy ranges.

In **Fig. 2C**, we present one example of a CPEB3 HDV-like ribozyme from Chimpanzee (PDB ID: 7QR3), a 69-nucleotide RNA molecule, where DRfold2 shows significant advantage in modeling tertiary structure. Among the four better-performing methods in our benchmark, DRfold2 accurately captures the topology of the ribozyme with a TM-score and RMSD of 0.586 and 2.77 Å, respectively. In contrast, DeepFoldRNA, while capturing the general helical arrangement, shows notable deviations in the orientation of the hairpin loop. As a result, the RMSD becomes 5.68 Å, which is twice the deviation of DRfold2. RhoFold and RoseTTAFoldNA show more significant errors, particularly in predicting the spatial organization of the junction regions, with TM-scores reducing to 0.323 and 0.285 respectively. Note that the maximum sequence identity between this target and the training RNAs is only 60.9%, demonstrating that the DRfold2 pipeline can provide reasonable structure predictions even for novel sequences that are not homologous to the training data.

### DRfold2 networks enhance the accuracy of RNA secondary structure prediction

Many RNA functions are closely associated with their secondary structure arrangements, which are primarily stabilized by hydrogen bonding and base-pair stacking interactions ^33^. **Figs. 3A-C** analyzes the Interaction Network Fidelity (INF) indices, which are defined as the geometric mean of precision and recall of secondary structure predictions, i.e., 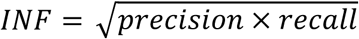. First, for Watson-Crick base pairs (INF_wc) and stacking interactions (INF_stack), while most methods (except for trRosettaRNA) demonstrate reasonable performance, DRfold2 achieved the highest INF values of 0.836 and 0.732. For non-canonical interactions (INF_nwc), however, all methods, including DRfold2, exhibit much lower INF values. Although DRfold still achieves the highest INF_nwc, its low value (0.176) underscores that accurately modeling non-canonical base-pair interactions remains an urgent challenge in RNA structure prediction. **Fig. 3D** presents a combined secondary structure accuracy across all three indices, where DRfold2 achieves INF_all=0.740, which is 5.1% higher than the second-best method, DeepFoldRNA (INF_all = 0.704).

**Figure 3.**
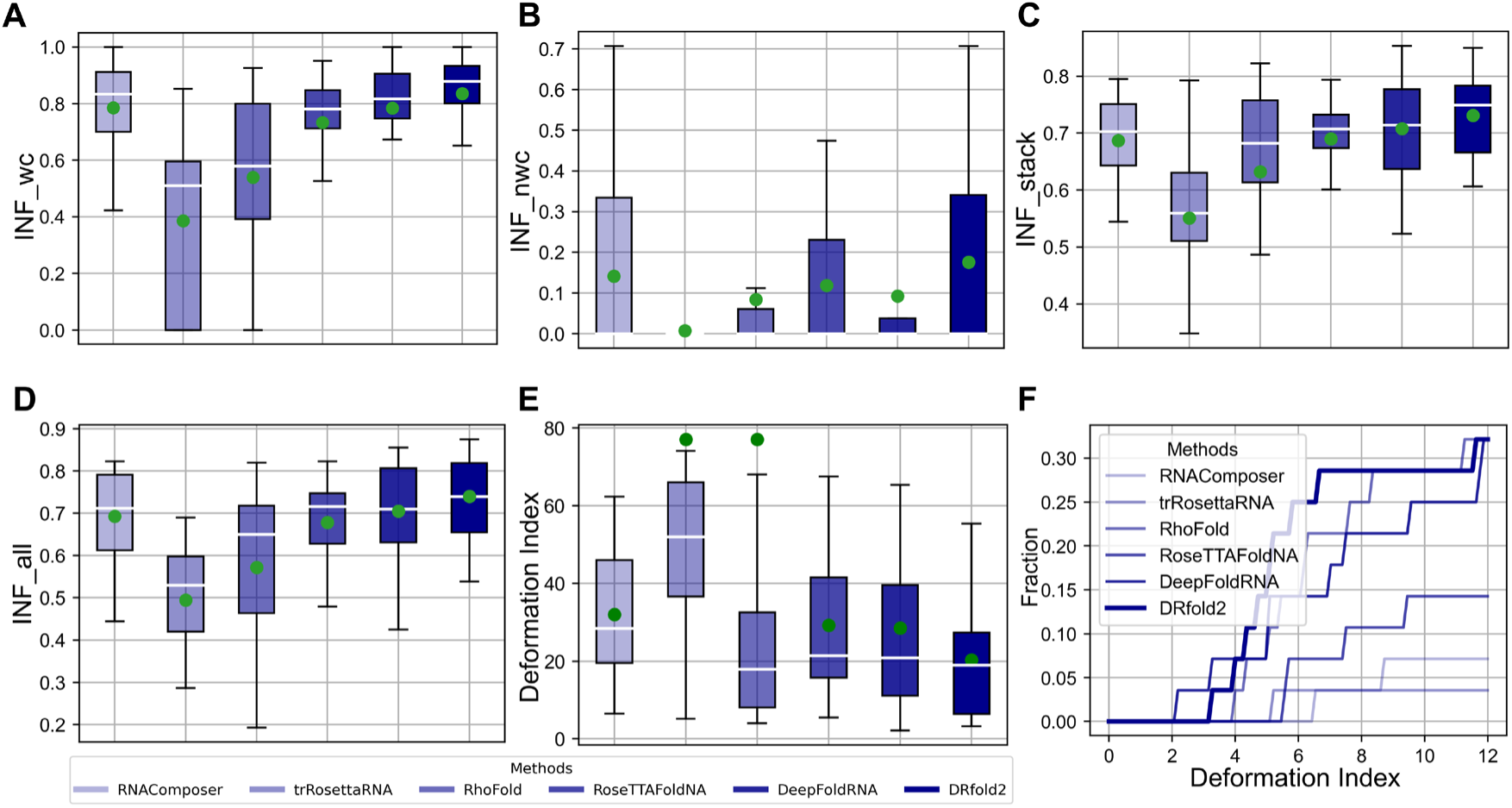
Comparative analysis of RNA secondary structure prediction. **(A-E)** Box plots comparing six RNA structure prediction methods across different RNA secondary structure performance metrics. The green point and the white horizontal line represent the mean and median values respectively. Note that the mean Deformation Indexes for trRosettaRNA and RhoFold are significantly higher than the median, due to their poor performance on a few targets where no correctly modeled base pairs are present in the predicted structures. **(F)** Cumulative distribution plot of deformation index for different methods.

While the INF indices assesses the conformational accuracy at the nucleotide pair level, the Deformation Index (DI), defined as *RMSD/INF_all*, has been introduced to provide a more balanced evaluation of 3D topological and 2D base-pairing patterns^34^. As shown in **Fig. 3E**, DRfold2 achieves the lowest average DI of 20.3, which is 40% lower than the second-best method, DeepFoldRNA (DI=28.5), where the corresponding p-value of 8.19E-03 indicates that this advantage is statistically significant. **Fig. 3F** further presents the accumulative fraction distribution at different DI cut-offs, where DRfold2 outperforms all control methods, achieving the highest area under the curve (AUC=2.04), which is 15% higher than the best control method, RhoFold (AUC=1.78). These results suggest that DRfold2 excels not only in predicting global structural topology but also in capturing detailed hydrogen-bonding and base-pair stacking patterns within RNA structures.

It is notable that DRfold2 has achieved superior secondary structure predictions using sequence information alone, without incorporating any precomputed secondary structure predictions, as many control methods do. This success can be attributed to several key components of DRfold2. First, the RCLM effectively captures inter-residue co-evolutionary patterns, providing reliable embedding features for the deep learning model. Second, the Denoising Structure Module helps enhance the end-to-end learning of inter-nucleotide frames through supervised training, ensuring proper spatial relationships between nucleotides. Finally, the precise prediction of inter-nucleotide geometries through multi-task learning helps guide the refinement of base pairing conformations during the post-processing optimization stage.

### RCLM more effectively captures co-evolutionary patterns than existing language models

One of the main innovations of the DRfold2 pipeline is the introduction of RCLM which is expected to provide a more precise approximation of the full likelihood probability by considering higher-order inter-nucleotide interactions (see **Methods**). To examine the assumption, **Figs. 4A-B** evaluate the performance of RCLM against RNA-FM, a representative RNA sequence language model^35^ trained on RNAcentral database^36^ with conventional Masked Language Model Loss. Two sequence-level metrics, i.e., sequence recovery rate and pseudo perplexity, calculated by sequentially masking each position in test sequences, have been used for assessment. Here, perplexity serves as a metric to evaluate the performance of a probabilistic model in masked sequence prediction, quantifying the language model’s uncertainty, where lower perplexity indicates higher confidence and better predictive accuracy^37^. The data show an improved masked nucleotide recovery ability for RCLM compared to RNA-FM. For instance, the average recovery rate and the perplexity of RCLM are 46.7% and 1.164 respectively, compared to 42.3% and 1.210 by RNA-FM. In **Fig. 4C**, we further analyzed the nucleotide-level perplexity across all 2,997 nucleotides in the 28 test RNAs, where RCLM again shows significantly better performance with an average perplexity of 1.186 versus 1.221 of RNA-FM (p-value = 5.19E-05 in one-sided Student’s t-test). Notably, these evaluation metrics are marginalized across positions, representing only a subset of RCLM. Even with this partial probability, RCLM demonstrates consistently superior performance over RNA-FM.

**Figure 4.**
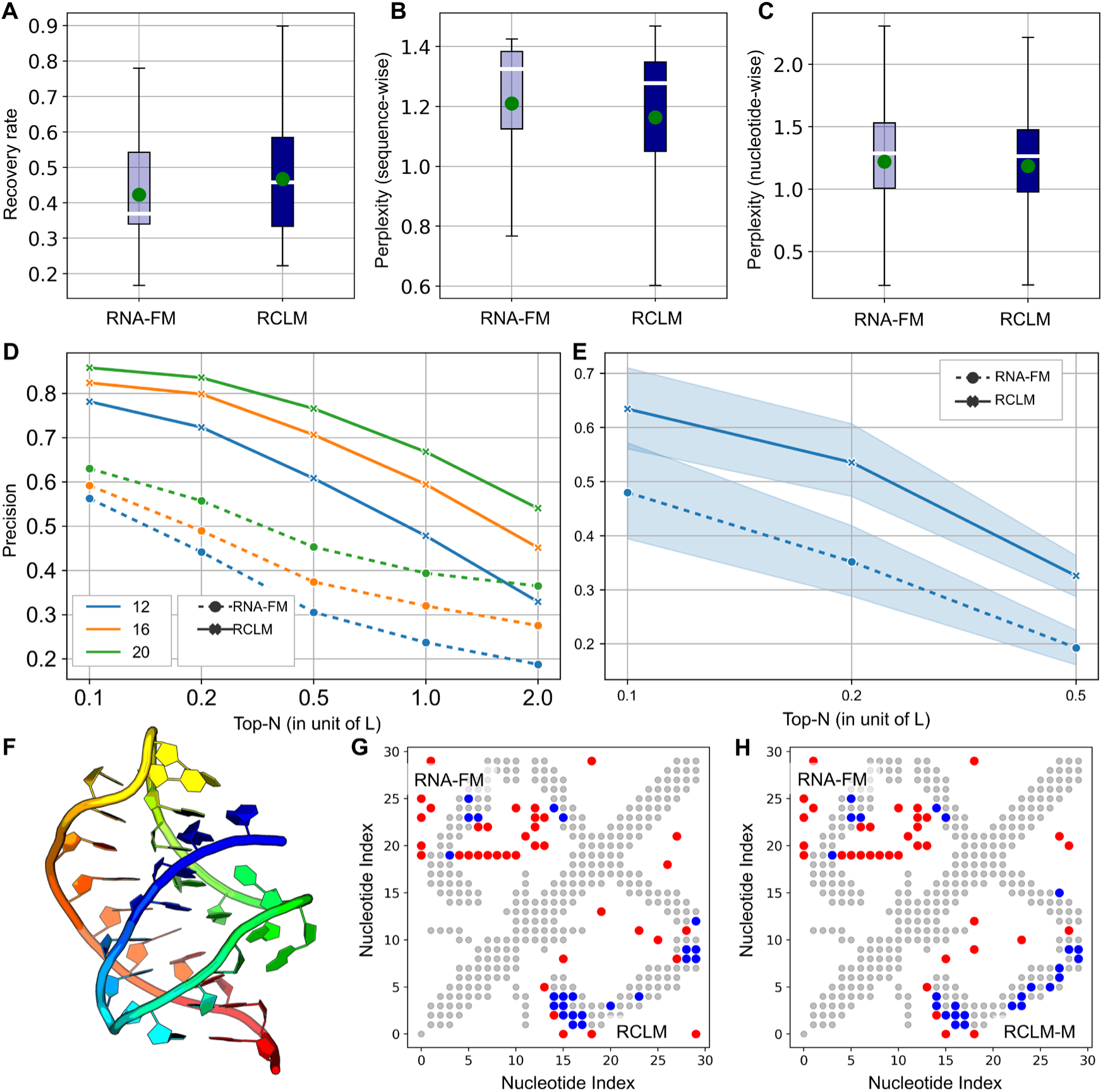
Comparative analysis of RCLM and RNA-FM for RNA sequence learning. (A-C) Box plots of sequence recovery rate, sequence-wise perplexity, and nucleotide-wise perplexity by RNA-FM and RCLM, respectively. Green points indicate means and white horizontal lines show medians. **(D)** Top-*N* precision curves at different contact thresholds (12, 16, and 20 Å) for RCLM and RNA-FM models. **(E)** Precision comparison of unsupervised RNA secondary prediction between RNA-FM and RCLM. **(F)** 3D structure visualization of the Class I type III preQ1 riboswitch from E. coli (PDB ID: 8FZA). **(G-H)** Contact maps of 8FZA, with grey dots representing ground-truth contacts, red dots false predicted contacts, and blue dots true positive predictions. Upper-left triangle shows results of RNA-FM predictions, and lower-right triangle is that of RCLM (**G**) and RCLM-M (**H**) predictions.

During the training, the composite likelihood of RCLM was calculated based on the sum of two probabilities for nucleotide-wise masked type and pair-wise joint predictions. In **Fig. S1A-C**, we also examine the sequence recovery ability derived from the pair-wise joint probability. Here, we marginalize the predicted joint distribution into nucleotide-wise distribution for evaluation, denoted as RCLM-M. Interestingly, the marginalized prediction achieves slightly better performance, with a sequence recovery rate of 48.7% and sequence- and nucleotide-wise perplexities of 1.149 and 1.175, compared to those with the direct prediction by RCLM (46.7%, 1.164 and 1.186), respectively. The result suggests that RCLM generates meaningful predictions for the pair-wise joint nucleotide distributions.

In addition to perplexities, a more critical assessment is on the co-evolution patterns captured by the language models, which are directly relevant to 3D RNA structure predictions. To evaluate this capability, we employed a completely unsupervised method, Categorical Jacobian ^38^, to extract the RNA contact maps from the language models. **Fig. 4D** presents a comparison of top-*N* contact map precision between RNA-FM and RCLM across multiple distance thresholds, where RCLM consistently has dramatically higher precisions than RNA-FM for all contact thresholds. For example, when using a contact distance cutoff of 12 Å, the Top-*L* precision of RCLM (47.8%) is more than 100% higher than that achieved by RNA-FM (23.7%). In **Fig. 4E**, we also compare the precision of unsupervised secondary structure prediction, by assuming that the Categorical Jacobian produces the base pairing score maps, where RCLM shows again a significant advantage over RNA-FM. These results indicate that RCLM can more effectively learn high-quality co-evolution patterns from sequence data than existing language modeling approaches. Since Categorical Jacobian requires mutation of all positions into all possible nucleotides during the extraction process, the high contact precision by RCLM also suggests that the composite language model has achieved a reliable level of sensitivity. In **Figs. S1D-E**, we further examine the precisions of the Categorical Jacobian contact and secondary structure predictions based on marginalized pairwise prediction from RCLM. While RCLM-M produces slightly lower precisions than RCLM, it still maintains a large performance margin compared to RNA-FM. It is important to note that RCLM achieves this superior performance with just 50% of the model size (47.50 million parameters for RCLM compared to 99.52 million for RNA-FM), further highlighting RCLM’s high efficiency in sequence language model learning.

As an illustration, **Figs. 4F-H** showcase an example of a Class I type III preQ1 riboswitch from E. coli (PDB ID: 8FZA) ^39^, which consists of 30 nucleotides as shown in **Fig. 4F**. The upper-left and lower-right panels of **Figs. 4G-H** are the unsupervised Top-*L* predicted contacts by RNA-FM and RCLM/RCLM-M respectively, where the ground-truth contacts (defined by inter-N atom distances < 12 Å) are shown as grey dots. RNA-FM only predicts 6 out of 30 (20%) positive contacts, while RCLM and RCLM-M correctly produce 17 (56.67%) and 19 (63.33%) positive contacts when using single nucleotide prediction and marginalized pairwise prediction, respectively, both exceeding RNA-FM’s accuracy by more than twofold. The result of this example highlights again the efficiency of RCLM to learn co-evolution information that is crucial for producing higher-precision structural restraints and *ab initio* RNA models, especially for the cases with limited sequence and structural homologies.

### Positive impact of composite language model on RNA 3D structure prediction

To directly examine the effectiveness and impact of different language models on 3D RNA structure prediction, we present in **Fig. 5A** the TM-scores of identical end-to-end structural learning networks built on one-hot, RNA-FM, and RCLM representations. It is observed that models using RCLM features achieve the highest mean and median TM-scores of 0.326 and 0.296, which are 7.9% and 10.9% higher than that with RNA-FM (0.302 and 0.267) and 10.1% and 16.5% higher than that of the one-hot representation (0.296 and 0.254), respectively. **Figs. 5B** and **5C** further list head-to-head TM-score comparisons of RCLM against the control models, where RCLM outperforms one-hot and RNA-FM in 20 and 17 cases, while underperforming in only 8 and 11 cases, respectively. To quantitatively examine the contribution of language models to the 3D RNA structure modeling, **Fig. 5D** shows the relationship between the TM-score of RCLM-based models and the Top-*L*/5 secondary structure precision from RCLM Categorical Jacobian, where a high Pearson Correlation Coefficient (PCC=0.528) is observed, suggesting that a higher RCLM-based secondary structure precision generally leads to more accurate 3D structure prediction. In addition to base pairing precision, **Fig. 5E** also examines the correlation between unsupervised contact map precision and TM-score, which shows a slightly weaker but robust PCC of 0.321; moreover, if we compare the improved TM-score (defined as the difference between RCLM-based and one-hot-based predictions), the PCC increases to 0.390 (**Fig. 5F**). Collectively, these results suggest that the comprehensive probability representations provided by advanced language models, like RCLM, significantly enhance the ability to learn co-evolutionary patterns and spatial restraints, leading to more accurate 3D RNA structure modeling through the end-to-end network in DRfold2.

**Figure 5.**
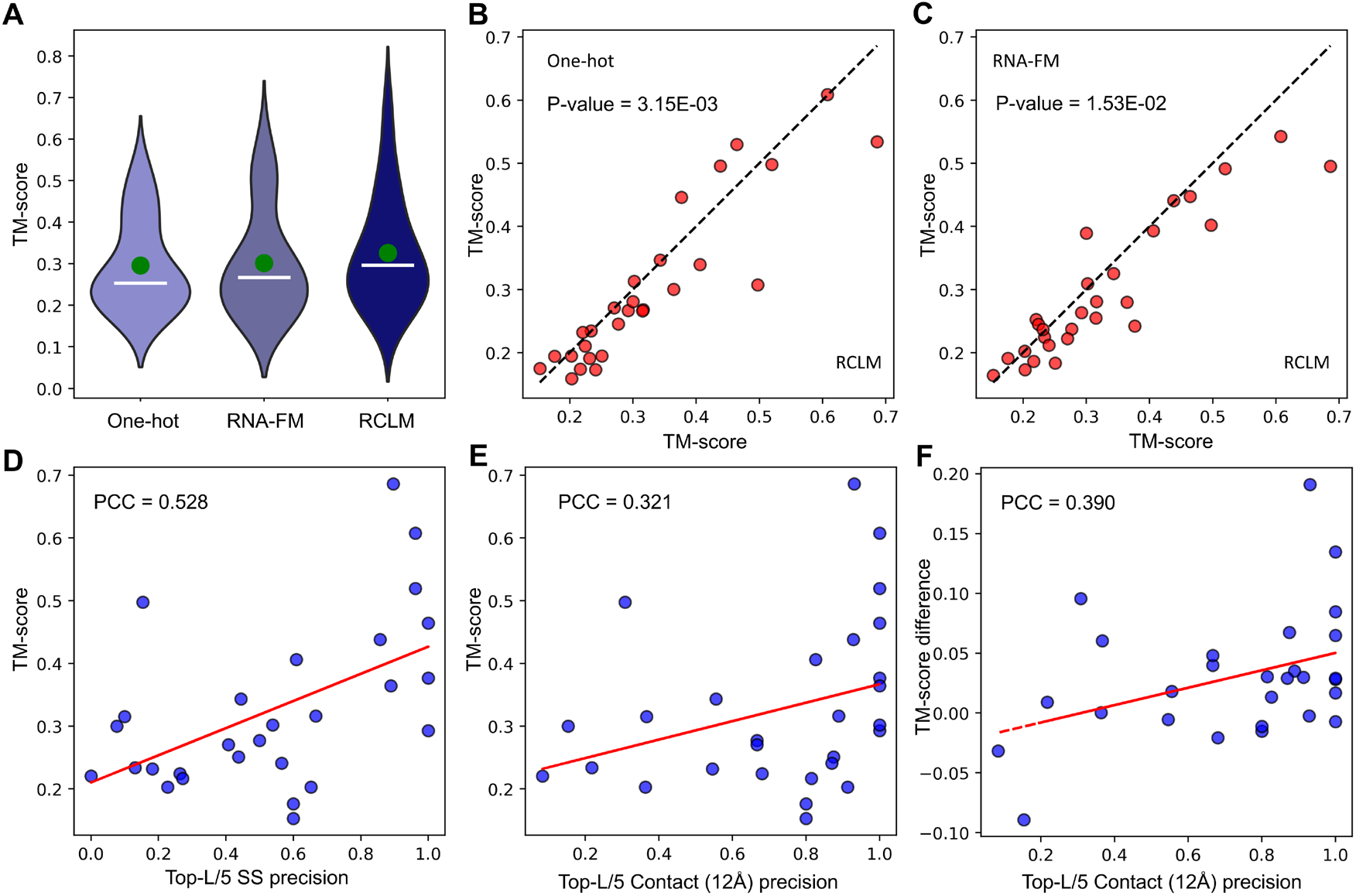
Comparison of RNA structure prediction performance across different encoding methods and correlation analysis between unsupervised contact/secondary structure precision and TM-score. **(A)** Violin plots of TM-scores by end-to-end RNA structure predictions based on different encoding strategies (One-hot, RNA-FM, and RCLM), with white horizontal lines indicating medians and green points showing means. **(B-C)** Head-to-head TM-score comparisons between RCLM and One-hot and RNA-FM encoding based models. **(D)** Correlation between TM-score and Top-L/5 secondary structure precision. **(E)** Correlation between TM-score and Top-L/5 Contact precision at 12Å threshold. **(F)** Correlation between TM-score difference and Top-L/5 Contact precision at 12Å threshold. Red lines indicate linear regression fits.

### DRfold2 is complementary with AlphaFold3

A comprehensive model of AlphaFold, AlphaFold3, was recently released^40^, and it is therefore of interest to compare the performance of DRfold2 and AlphaFold3 for RNA 3D structure prediction. For doing this, we submitted the RNA sequences in our test set to the AlphaFold Server and obtained the predicted structures of AlphaFold3 with default seed configurations. It was found that on average, DRfold2 achieves a slightly higher TM-score (0.351) and lower RMSD (14.6 Å) compared to AlphaFold3 (0.345 and 16.0 Å) (**Table S1**). **Figs. 6A-B** provide head-to-head TM-score and RMSD comparisons by the two programs. Although DFfold2 outperforms AlphaFold3 in several low- to medium-quality models, AlphaFold3 excels in two targets with TM-score above 0.6.

**Figure 6.**
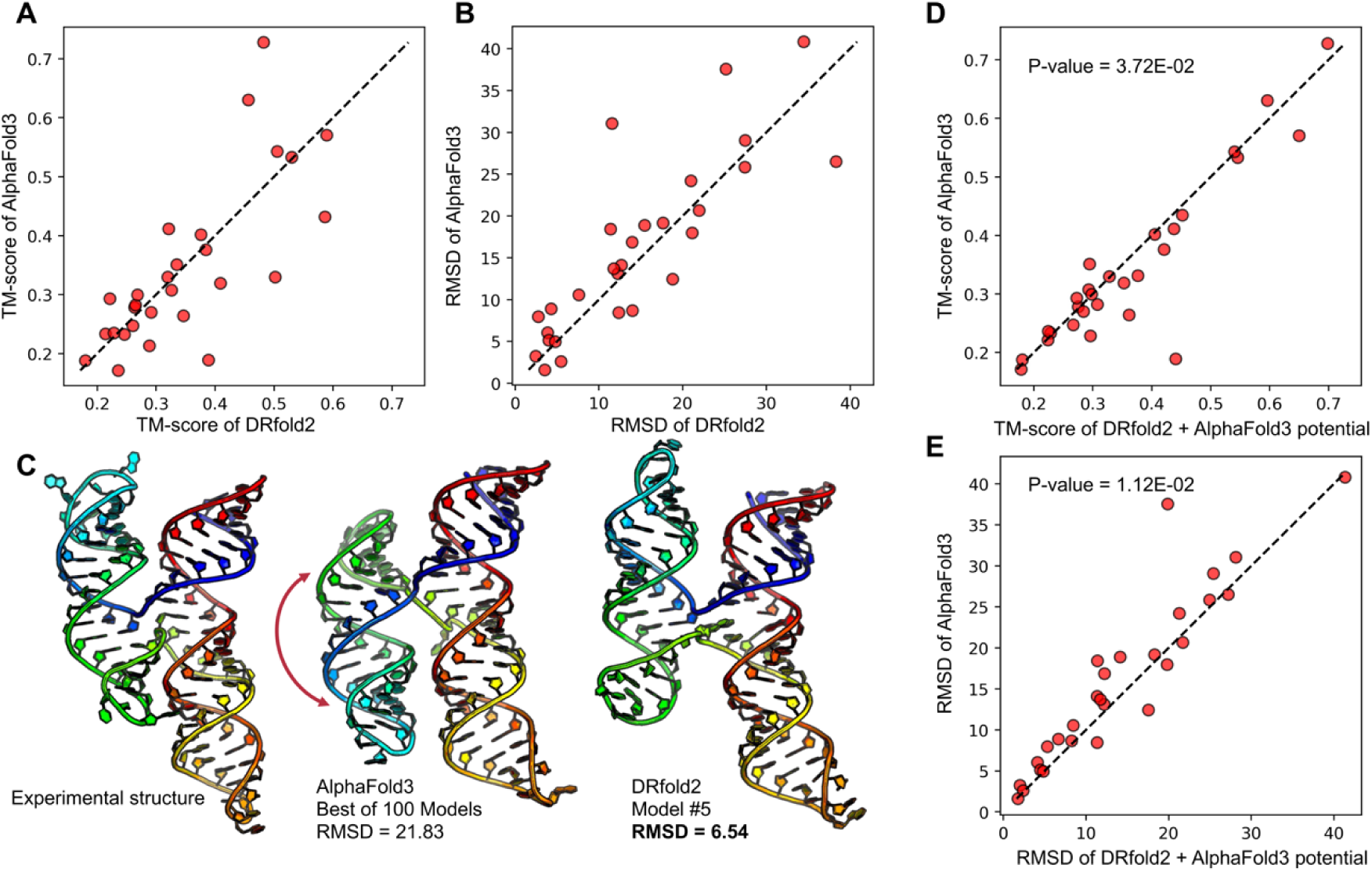
Comparative analysis of DRfold2 and AlphaFold3 for RNA structure predictions. (A-B) Head-to-head TM-score and RMSD comparisons between DRfold2 and AlphaFold3. **(C)** Structural visualization of the example from coxsackievirus B3 cloverleaf RNA (PDBID: 8DP3), showing experimental structure, AlphaFold3’s best prediction from 100 models, and 5^th^ model of DRfold2, respectively. Structures are rainbow-colored from 5’ (blue) to 3’ (red) end. **(D-E)** TM-score and RMSD comparisons between AlphaFold3 and DRfold2 with DRfold2+AlphaFold3 potential.

While most of our above discussions focus on the top-rank models by DRfold2, we have identified several targets for which DRfold2 generated correct structures, but the post-process step failed to select them as the first-rank models. **Fig. 6C** illustrates one such examples from the coxsackievirus B3 cloverleaf RNA (PDB ID: 8DP3), which contains 90 residue forming a 4-way junction fold ^41^. By checking the first-rank models, both DRfold2 and AlphaFold3 failed to correctly predict the direction of the 4-way junction, with RMSDs to the native around 20 Å. To further explore AlphaFold3’s capabilities, we submitted the same sequence to AlphaFold Server 20 times with different seeds, resulting in 100 predicted conformations. The middle panel of **Fig. 6C** shows the best conformation with the highest TM-score, which still exhibits a poor RMSD of 21.83 Å due to incorrect chain winding topology across the 4-way junction. In contrast, the 5^th^-ranked model from DRfold2 successfully predicted the junction’s direction, significantly reducing the RMSD from 21.96 Å (first DRfold2 model) to 6.54 Å (right panel of **Fig. 6C**). While this example highlights DRfold2’s capability to generate diverse and alternative conformations, it also underscores the need to improve the post-processing SCOR protocol for more effectively selecting and ranking the correct models from the DRfold2 decoy pool, e.g., through the combination of third-party algorithms.

Although DRfold2 and AlphaFold3 exhibit similar overall performance, the results in **Figs. 6A-B** highlight the strong complementarity between the two programs, particularly in cases where predictions deviate significantly from the diagonal line. This complementary behavior, along with the identified need to improve DRfold2’s model selection strategy, motivated us to explore whether combining the two approaches could lead to enhanced structure predictions. The optimization module in DRfold2 readily facilitates this integration by incorporating AlphaFold3 conformations as an additional potential term, denoted as DRfold2+AlphaFold3 potential. Guided with this combined potential function, the optimized structures achieve higher quality models than those generated by either method alone. For example, as compared in **Figs. 6D-E**, the hybrid approach achieves an average TM-score of 0.365 and RMSD of 14.21 Å, both being significantly better than that of AlphaFold3 (0.345 and 16.022 Å, with p-value = 3.72E-02 for TM-score and 1.12E-02 for RMSD). These results not only validate our hypothesis regarding the complementarity between DRfold2 and AlphaFold3 but also demonstrate the flexibility and extensibility of the SCOR module, enabling the seamless integration of external model features to further improve structure prediction in DRfold2.

### Performance of DRfold2 in CASP16

An earlier version of DRfold2 participated in the 16^th^ Critical Assessment of protein Structure Prediction (CASP16) experiment for RNA structure prediction under the group ID ‘dNAfold’. **Table S2** summarizes the average TM-score and accumulated Z-scores for the Top 20 groups on the 20 RNA targets with known experimental structure and sequence lengths below 400 nucleotides. Although the CASP16 assessment contains both RNA and DNA targets, only monomer RNA targets, for which dNAfold submitted predictions, were included in this analysis.

The results revealed that Vfold, which utilizes a template-based modeling approach integrated with AlphaFold3^42^, achieved the highest performance, with an average TM-score of 0.566 and Z-score of 19.71, significantly outperforming other groups. The remaining top-ranked groups had comparable average TM-scores around 0.5, with DRfold2 securing 5^th^ place in both TM-score (0.510) and Z-score (13.46), indicating its strong competitiveness among the state-of-the-art programs. Consistent with the benchmark results shown in **Fig. 6**, DRfold2 slightly outperformed the AlphaFold3 server, which was ranked 7^th^ in TM-score (0.501) and 16^th^ in Z-score (9.78) (**Table S2**).

It is noteworthy that the AlphaFold3 server was released before the CASP16 experiment, allowing the participating groups to integrate AlphaFold3 predictions into their modeling pipelines, which many did^43^. However, we chose not to incorporate AlphaFold3 or any third-party predictions in our CASP16 submissions, aiming to more critically assess the DRfold2’s independent modeling capabilities. To further assess the uniqueness of the RNA structure predictions, we introduced an AlphaFold dissimilarity score, defined as 1 − *TM_M_*, where *TM_M_* is the maximum TM-score between the first submitted model by each group and the five models submitted by AlphaFold3 (Group ID: AF3-server). In **Fig. S2**, we plot the average TM-score on 20 RNA monomer targets versus the AlphaFold dissimilarity for all groups that submitted predictions for all these targets, where dNAfold exhibited the highest dissimilarity score among the six groups that have an average TM-score outperforming AlphaFold3. Moreover, dNAfold achieved the top rank among all groups with a dissimilarity score above 0.45. These results highlight the capability of DRfold2 for producing distinct and independent structure predictions. Nevertheless, given the complementary nature of DRfold2 and AlphaFold3 in both methodology and the benchmark results shown in **Fig. 6**, we expect that appropriate integration of DRfold2 with AlphaFold3 or other top-performing models represents a promising direction for future improvements in DRfold2-based RNA structure prediction.

## Discussion

RNA structure prediction has been largely constrained by the limited availabilities of experimental structures that can be used for template recognitions and/or training effective AI models. We have developed DRfold2, a new deep learning-based method for *ab initio* RNA structure prediction by integrating a newly trained RNA Composite Language Model with a denoising-coupled end-to-end learning strategy. The method has been tested on a non-redundant test set with different redundancy cut-offs and shown consistently improved performance over existing approaches in both 3D tertiary structure prediction and 2D base pair modelling, demonstrating the effectiveness and robustness of the proposed method.

The key factor contributing to the success of DRfold2 is the proposed RCLM, trained from scratch with an improved loss function for better likelihood approximation. The RCLM model achieves significantly higher unsupervised contact prediction precision compared to existing representative methods, demonstrating a superior ability to capture co-evolutionary signals solely from sequence data. The correlation between contact precision and structural accuracy further validates its effectiveness in learning structurally relevant patterns directly from sequences. Meanwhile, the newly introduced Denoising RNA Structural Module (DRSM) enhances both the efficiency and robustness of the end-to-end training process.

While DRfold2 shows comparable overall performance to AlphaFold3 on our test set, our analysis suggests that the two methods provide highly complementary predictions. By incorporating AlphaFold3 predictions as an additional potential term in DRfold2 optimization framework, we achieved statistically significant improvements in both TM-score and RMSD, compared to either approach.

The CASP16 results further validate the effectiveness of DRfold2, particularly for regular-sized monomer RNAs. However, we observed relatively lower performance and ranking on large RNA targets above 200 nucleotides. One possible reason is that many groups incorporated AlphaFold3, which has twice the crop size of DRfold2, whereas we intentionally evaluated DRfold2 independently, relying solely on our own method for testing. We understand that the CASP competition naturally emphasizes prediction accuracy of submitted models, which often leads participants to combine multiple existing methods for the best possible results. However, developing new independent methods remains important for the field. DRfold2 offers a distinct approach that uses the composite language model, deep end-to-end learning, and post optimization protocols, providing a different perspective on RNA structure prediction, as highlighted in the AlphaFold3 dissimilarity analysis by **Fig. S2**. Although it may not always outperform top-performed models like AlphaFold3, the diversity in modeling approaches prevents over-dependence on a single method and creates opportunities for future innovations.

## Materials and methods

DRfold2 is new pipeline for RNA 3D structure prediction consisting of four core modules: (1) RNA Composite Language Model, (2) RNA Transformer Block, (3) Denoising Structure Module, and (4) final model selection and refinement through the CSOR Protocol (**Fig. 1**). While it shares a similar name with the earlier DRfold method^21^, DRfold2 introduces significant advancements built on a completely different framework, most notably with the incorporation of the Composite Language Model, which greatly enhances RNA sequence and pairwise representations. Additionally, the pipeline integrates a Denoising RNA Structural Module (DRSM), which employs a controlled perturbation strategy to robustly learn structural transformations by efficiently correcting noisy RNA conformations. The pipeline also features a carefully designed flexible end-to-end loss function with distance-dependent reweighting schema to help the model learn meaningful interaction patterns. Lastly, we extend the deep learning-based conformational pool for the proposed CSOR to identify the most promising structure and possible alternative conformations. Equipped with those advancements, DRfold2 significantly outperforms DRfold (**Fig. S3**), providing more than three times as many targets (from 2 to 7) foldable RNAs (TM-score > 0.45) on the test set used in this study.

### RNA sequence representation learning by Composite Language Model

The first component of DRfold2 is a pre-trained RNA language model, RCLM, which can provide enhanced RNA sequence representation. Given an input RNA sequence with *L* nucleotides, which is represented by one-hot encoding, two types of sequence and pair representations will be generated through RCLM (**Fig. 1B**). The initial sequential representation 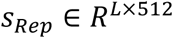 can be obtained by applying a linear layer over the one-hot feature. In addition, another two separate sequence projections above *S_Rep_* will be added vertically and horizontally to form the initial pair embedding 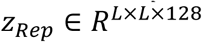. Here, 512 and 128 are the representative dimensions for sequence and pair features, respectively.

The initial sequential and pair representations, *S_Rep_* and *Z_Rep_*, will be used as the input of a set (18 in RCLM) of RNA Transformer Blocks. As shown in **Fig. 1C**, each transformer block is designed to model pairwise interactions, which are critical for biomolecules. It is worth noting that two sub-modules, i.e., Triangle Attention StartingNode and Triangle Attention EndingNode modules, were omitted in RCLM, as this configuration significantly reduces the GPU memory consumption.

In addition to the pair modeling, a key innovation of RCLM is the implementation of composite likelihood maximization for masked regions during the training. Given a sequence *x* = (*x*_1_, *x*_2_, ⋯, *x_L_*), the objective of masked language models is to maximize the probability of masked regions ℳ, i.e., *P*(*X*_*i*_ = *x*_*i*_, ∀_*i*_ ∈ ℳ). For notational simplicity, we omit the conditioning on unmasked regions and network parameters, though they are included in the full conditional probability. We find that existing mainstream BERT-based language models^44^ approximate this distribution by maximizing a *pseudo likelihood* ∏_*i*ɛℳ_ *P*(*X*_*i*_ = *x*_*i*_), assuming that each masked token *X*_*i*_ is independent of the others. However, this independence assumption may not always hold, especially for RNAs, where multiple interactions occur among nucleotides.

To better approximate the actual full likelihood in RNAs, we consider maximizing a composite likelihood (*CL*)^26^, given by

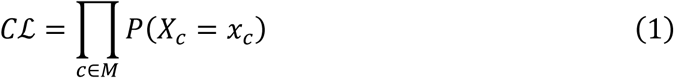

where *C* is a set of masked position subsets. When *C* = {{i: i ∈ ℳ}_, *C*ℒ corresponds to the full likelihood that models the joint distribution of all masked positions simultaneously. On the other hand, if we set *C*= {{i}, i ∈ ℳ}, *C*ℒ reduces to the pseudo likelihood, where each masked token is assumed to be conditionally independent of others. In our implementation of RCLM, we incorporate second-order interactions by setting *C* = {{i}, i ∈ ℳ } ∪ {{*i*, *j*}, *i* ∈ ℳ *j* ∈ ℳ}, which considers both individual tokens and interactions between pairs of masked positions. Thus, the final loss function for our RCLM training is the negative log-composite likelihood:

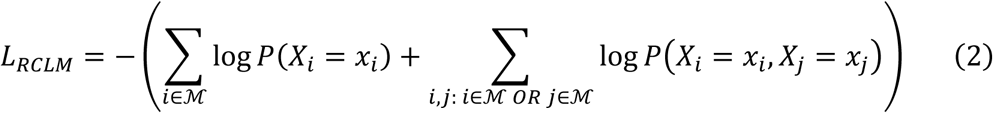

In this case, we consider both first- and second-order interactions, which naturally align with the sequential and pair representations learned by the RNA Transformer Blocks. Notably, composite likelihood maximization has been previously applied in protein sequence modeling, such as clmDCA^45^ for direct coupling analysis. Here, we incorporated pairwise composite likelihood maximization into RNA language model training, to enable the capture of complex nucleotide interactions that go beyond simple positional independence assumptions.

The RCLM was trained on ∼30M RNA sequences from the RNAcentral database^36^ (Release 22) over 67,000 batches, with a batch size of 128. The entire training process took about 15 days using a single NVIDIA A40 GPU.

### RNA structure representation learning with RNA Transformer Block

After training RCLM, the sequence and pair representations obtained from its final layer are utilized as the input for RNA structure prediction. The backbone framework is the complete RNA Transformer Block, as illustrated in **Fig. 1C**. The input sequence representation is first processed through an Attention module, where a bias term, derived from the linear projection of the input pair representation, is added to the attention weights. This bias can effectively incorporate pairwise information to the sequence modeling. The resulting sequence representation is then passed through a Transition Module to generate the final output sequence representation.

A separate copy of the output sequence representation services as the input of a set of pair processing modules, including Triangle Multiplication Outgoing, Triangle Multiplication Incoming, Triangle Attention StartingNode, Triangle Attention EndingNode, and a final Pair Transition Module. In our implementation, the dimensions of sequence and pair representations are uniformly set to 64. For sequence representation learning, we set the number of attention heads to 8, with each head having a channel size of 8. Due to the high memory requirements of pair-related modules, however, we reduce the number of heads to 4. After passing through 16 RNA Transformer Blocks, we obtain the sequence and pair representations, *s* ∈ 𝑅^𝐿×64^ and 𝑧 ∈ 𝑅^𝐿×𝐿×64^, which will be used for subsequent structure prediction.

During sequence-to-structure representation learning in DRfold2 through RNA Transformer Blocks, we introduce an additional attention dropout in regular attention operations. We expect that such a technique will prevent the model from over-relying on specific input tokens in the context. Specifically, when updating the token representations, we randomly set 25% of the attention weights to zero in the attention map. To maintain stability, we normalize the masked attention map by scaling the remaining attention weights based on their sum along the weighting dimension. During inference, we disable the attention dropout operation, allowing the model to fully utilize the input context.

### End-to-end RNA structure prediction with the Denoising Structure Module

The sequence and pair representations obtained from the last RNA Transformer Block will be utilized as the inputs of the Denoising RNA Structure Module, as illustrated in **Fig. 1D**. Each input representation first passes through an independent LayerNorm layer. An Invariant Point Attention (IPA) Module ^20^ iteratively takes the normalized features as inputs to get nucleotide-wise features, which will be subsequently used to predict nucleotide-wise rotation matrices and translation vectors.

Another required input for the IPA Module is the initial structural conformation. We apply black-hole initialization ^20^, where each nucleotide’s initial state is defined by an identity rotation matrix 𝑅_0_ = 𝐼 and a zero-translation vector 𝑡_0_ = 0. During training, we employ a denoising strategy by adding controlled perturbations to these initial transformations. For rotation matrices, we apply weighted spherical interpolation *R_noised_* = *slerp*(*I*, *R_random_*, *w*) is a random rotation matrix, and 𝑤 = 0.1 is the interpolation weight. For translation vectors, we add random noise and *t_noised_* = 𝑡_0_ + 𝜀, where 𝜀∼ N(0, 0.1) denotes noise drawn from a normal distribution with mean 0 and scale 0.1. Given the high flexibility of RNA molecules, such controlled noise injection is expected to help improve model robustness during training and prediction.

Besides the initial rotation matrices and translation vector, the local nucleotide conformation 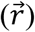 of three atoms, i.e., P, C4’, and glycosidic N atoms of the nucleobase, should also be pre-determined in the structure module. Following our previous approach^21^, symmetric orthogonalization^46^ is utilized to obtain the coordinates for each atom from the ideal A-form RNA helix structure. Such orthogonalization has proven to have only half of the reconstruction error compared to the Gram-Schmidt process used in AlphaFold models^47^.

In addition to the End-to-End structure models, DRfold2 also generates a set of inter-nucleotide distance predictions based on the pair representations from the last RNA Transformer Block, through a linear layer. Accordingly, the final loss function of DRfold2 for structure prediction consists of two terms:

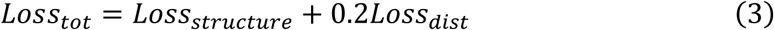

Where

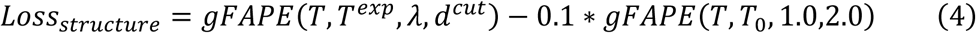

The first term in Eq. (4) quantifies the deviation between the predicted global nucleotide-wise rigid transformation (𝑇) and the experimental global nucleotide-wise rigid transformation (*T^exp^*), i.e.,

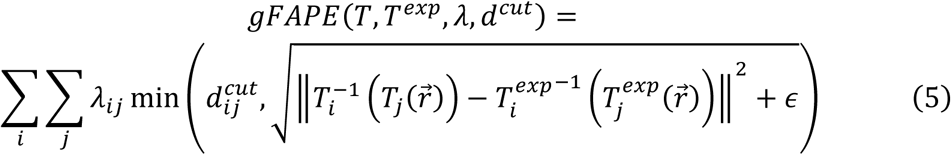

where 𝜆_*ij*_represents the inter-nucleotide re-weighting factor and 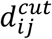 denotes the deviation cut-off parameter between nucleotides *i* and *j*. If the inter-nucleotide distance in the experimental structure is below 20 Å, we set 𝜆_*ij*_ = 3.0 and 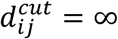. Otherwise, they are set to 1.0 and 30 Å, respectively. The value of 𝜖 is fixed at 1e-3. The subtraction of the second term in Eq. (4) encourages deviation from the initial black-hole transformation (𝑇_0_). Our experiments indicate that incorporating both the re-weighting scheme and the subtraction term accelerates the training process.

The second loss in Eq (3) for distance terms is defined as

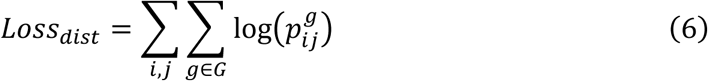

where 𝐺 includes the set of pairwise distances between P, C4’, and N atoms respectively for positions *i* and *j*. The distance values are evenly divided into 38 bins spanning 2-40 Å, with two additional bins for distances <2 and >40 Å. 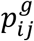 denotes the predicted probability in the respective distance bin 𝑔.

We trained four randomly initialized models for approximately 500,000 steps using a crop size of 256 tokens. The models were then trained for an additional 100,000 steps with a 384-token crop size. For each model, we retained the last 20 checkpoints saved at ∼2000-step intervals. This resulted in 80 checkpoints in total (4 models × 20 checkpoints each), which were used to create a conformational pool for further optimization.

### Structure optimization through CSOR protocol

During inference, for each input sequence, a collection of *N*=80 end-to-end structures and distance probability maps is generated, where the final predicted models are obtained through the CSOR protocol (Clustering, Scoring, Optimization and Refinement), as illustrated in **Fig. 1E**.

For **scoring**, we construct a global energy function *E_global_* which consists of two deep learning-based energy terms:

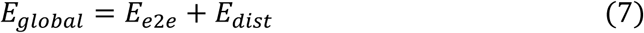

The first energy term incorporates the conformational information through gFAPE:

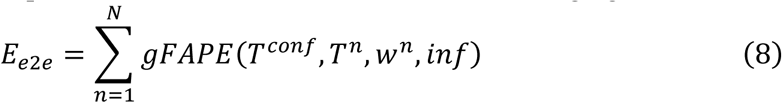

where *T^conf^* represents the transformations of the given conformation to be scored/optimized; 𝑇^𝑛^ is the predicted conformation of *n*-th out of 80 predicted structures; and 𝑤^𝑛^ is the predicted pairwise error estimation score of *n*-th checkpoint. The second term in Eq. (7) is defined by

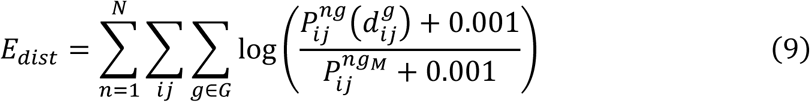

where 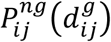 is the predicted probability of *n*-th prediction, given distance 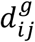 for the 𝑔-distance bin. 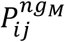 is the corresponding probability of the last distance bin (*g_M_*) below the upper threshold (40 Å).

Based on the lowest *E_global._*, we select the 5 predicted structures and their distance maps. A L-BFGS based **optimization** process ^48^ is performed under the guidance of *E_opt_* = *E_global_* but with *N* converted from 80 to 5 selected structures. The final conformation with the lowest *E_opt_* will be considered as the *first* predicted model.

In order to find possible alternative/variant conformations from our prediction pool, we performed a basic **clustering** step by SPICKER algorithm ^49^. Each cluster center will also go through the same scoring and optimization procedures described above. Eventually, up to 5 models will be obtained, and each of them will be **refined** by Arena ^50^ and OpenMM^51^.

## Data availability

The datasets collected and used in this work are available at https://zhanglab.comp.nus.edu.sg/DRfold2/data.zip. We show structures of 7QR3, 8FZA and 8DP3 obtained by four-digit accession codes in the PDB repository (https://www.rcsb.org/). The RNA sequence data for RCLM training was download from RNAcentral (https://rnacentral.org/).

## Code availability

The online server and standalone package of DRfold2 are freely available at https://zhanglab.comp.nus.edu.sg/DRfold2/. Data were analyzed using Numpy v.1.20.3 (https://github.com/numpy/numpy), SciPy v.1.7.1 (https://www.scipy.org/), Structures were visualized by Pymol v.3.0.0 (https://github.com/schrodinger/pymol-open-source)

## Acknowledgements

We thank Drs. Robin Pearce and Chengxin Zhang for discussions.

## Author contributions

Y.Z. conceived the project and designed the experiments; Y.L. developed methods and designed and performed experiments; C.F. prepared the initial data and performed the benchmark tests. X.Z. participated in the discussion and constructed the online server. Y.L. wrote the initial manuscript; all authors proofread and approved the final manuscript.

