## Supplementary Materials for "*Ab initio* RNA structure prediction with composite language model and denoised end-to-end learning"

#### Supplementary Tables

**Table S1.** TM-score and RMSD of the first model by DRfold2 and AlphaFold3 for all 28 test targets. Bold fonts highlight better performance in each subcategory.

| | TM-score | | RMSD (Å) | |
|---|---|---|---|---|
| Targets | DRfold2 | AlphaFold3 | DRfold2 | AlphaFold3 |
| 7elp_A | 0.329 | **0.412** | **3.87** | 6.05 |
| 7qr3_D | **0.586** | 0.432 | **2.77** | 7.98 |
| 7qr4_B | **0.502** | 0.330 | **4.30** | 8.89 |
| 7v9e_A | 0.320 | **0.330** | **12.24** | 13.10 |
| 7yr6_A | 0.263 | **0.278** | **27.45** | 29.06 |
| 7yr7_A | 0.214 | **0.233** | 27.42 | **25.85** |
| 8bu8_A | 0.221 | **0.293** | 38.29 | **26.52** |
| 8dp3_R | **0.409** | 0.319 | 21.96 | **20.66** |
| 8fza_A | **0.246** | 0.232 | **2.43** | 3.24 |
| 8gxc_A | **0.326** | 0.307 | 13.99 | **8.68** |
| 8hb8_A | **0.260** | 0.247 | **12.68** | 14.11 |
| 8hzd_A | 0.265 | **0.282** | **7.60** | 10.56 |
| 8hzl_A | **0.236** | 0.171 | **11.57** | 31.07 |
| 8its_A | **0.291** | 0.270 | **11.78** | 13.71 |
| 8jhp_A | **0.289** | 0.213 | **4.02** | 5.14 |
| 8qo2_A | 0.269 | **0.299** | **34.43** | 40.82 |
| 8qo3_A | **0.384** | 0.376 | **15.46** | 18.88 |
| 8s95_C | 0.530 | **0.533** | **13.97** | 16.86 |
| 8t2p_A | 0.335 | **0.351** | **17.65** | 19.17 |
| 8t2p_B | 0.376 | **0.402** | 21.13 | **17.98** |
| 8tvz_C | **0.388** | 0.189 | **25.16** | 37.55 |
| 8uo6_A | 0.505 | **0.543** | 18.79 | **12.45** |
| 8uo6_B | **0.589** | 0.571 | 12.37 | **8.46** |
| 8v1h_A | 0.457 | **0.631** | 5.46 | **2.62** |
| 8vci_A | 0.228 | **0.235** | **20.98** | 24.22 |
| 8vt5_A | 0.482 | **0.728** | 3.53 | **1.61** |
| 9bun_A | **0.346** | 0.264 | **11.40** | 18.42 |
| 9eow_A | 0.180 | **0.188** | **4.76** | 4.96 |
| Average | **0.351** | 0.345 | **14.552** | 16.022 |

**Table S2.** Average and Z-score based relative group performance of first model for TM-score for 20 RNA monomer targets (< 400 Nts) in CASP16. Only the top 20 groups are listed. Z-scores are calculated using a minimum Z-score cutoff >0.

| Average TM-score | | | SUM Z-score | | |
|---|---|---|---|---|---|
| Rank | Group ID | TM-score | Rank | Group ID | SUM Zscore |
| 1 | Vfold | 0.566 | 1 | Vfold | 19.71 |
| 2 | KiharaLab | 0.527 | 2 | GuangzhouRNA-human | 17.96 |
| 3 | CSSB_experimental | 0.520 | 3 | CSSB_experimental | 13.67 |
| 4 | Yang-Server | 0.516 | 4 | GuangzhouRNA-meta | 13.51 |
| **5** | **dNAfold** | **0.510** | **5** | **dNAfold** | **13.46** |
| 6 | elofsson | 0.505 | 6 | Yang-Multimer | 12.87 |
| **7** | **AF3-server** | **0.501** | 7 | KiharaLab | 12.82 |
| 8 | GuangzhouRNA-meta | 0.495 | 8 | GeneSilico | 12.49 |
| 9 | BRIQX | 0.493 | 9 | RNApolis | 12.34 |
| 10 | RNApolis | 0.493 | 10 | Yang-Server | 11.95 |
| 11 | Bhattacharya | 0.477 | 11 | BRIQX | 11.20 |
| 12 | Yang-Multimer | 0.473 | 12 | LCBio | 10.81 |
| 13 | RNAFOLDX | 0.466 | 13 | Diff | 10.60 |
| 14 | Diff | 0.442 | 14 | elofsson | 9.99 |
| 15 | Zheng | 0.434 | 15 | Bhattacharya | 9.92 |
| 16 | MIEnsembles-Server | 0.429 | **16** | **AF3-server** | **9.78** |
| 17 | NKRNA-s | 0.428 | 17 | GromihaLab | 9.71 |
| 18 | OpenComplex | 0.413 | 18 | RNAFOLDX | 7.55 |
| 19 | GuangzhouRNA_AI | 0.407 | 19 | CoDock | 7.19 |
| 20 | RNA_Dojo | 0.385 | 20 | falcon2 | 7.03 |

### Supplementary Figures

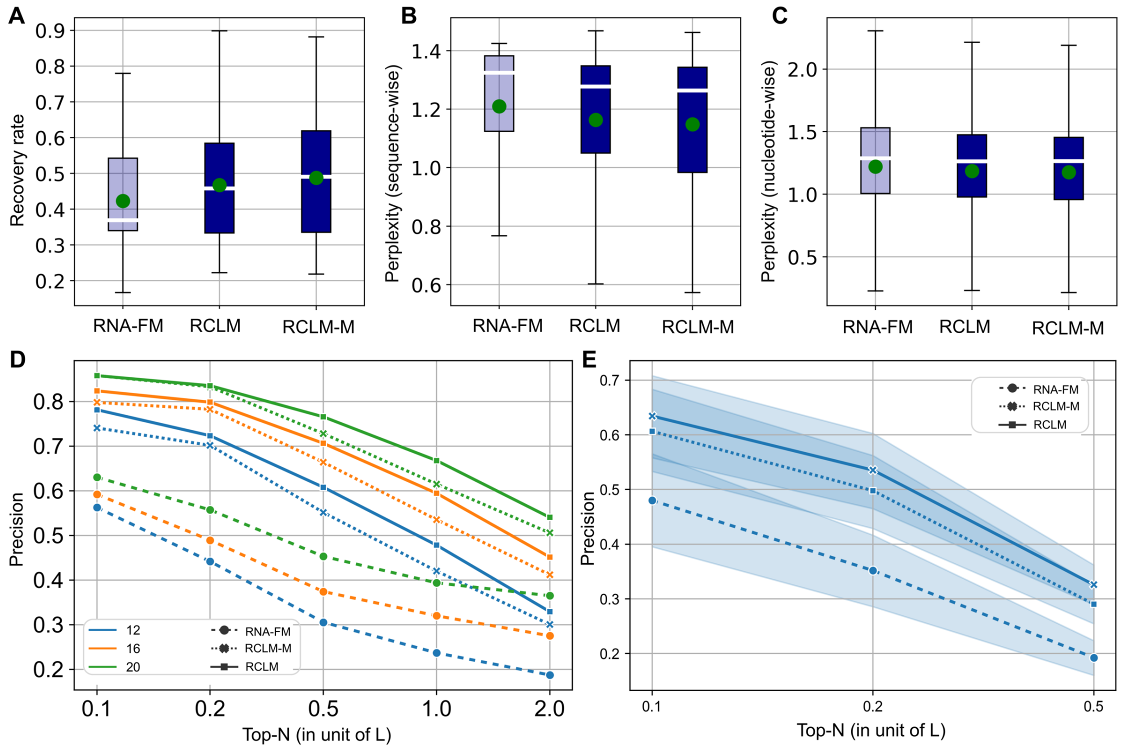


**Figure S1. Comparative analysis of RNA-FM, RCLM and RCLM-M methods for RNA sequence learning. (A-C)** Box plots comparing of sequence recovery rate, sequence-wise perplexity and nucleotide-wise perplexity between RNA-FM, RCLM and RCLM-M, respectively. Green points indicate means and white horizontal lines show medians. **(D)** Top-*N* Precision curves at different contact thresholds (12, 16, and 20 Å) for RNA-FM, RCLM and RCLM-M models, where *L* is the RNA sequence length. **(E)** Precision comparison of unsupervised RNA secondary prediction between RNA-FM, RCLM and RCLM-M.

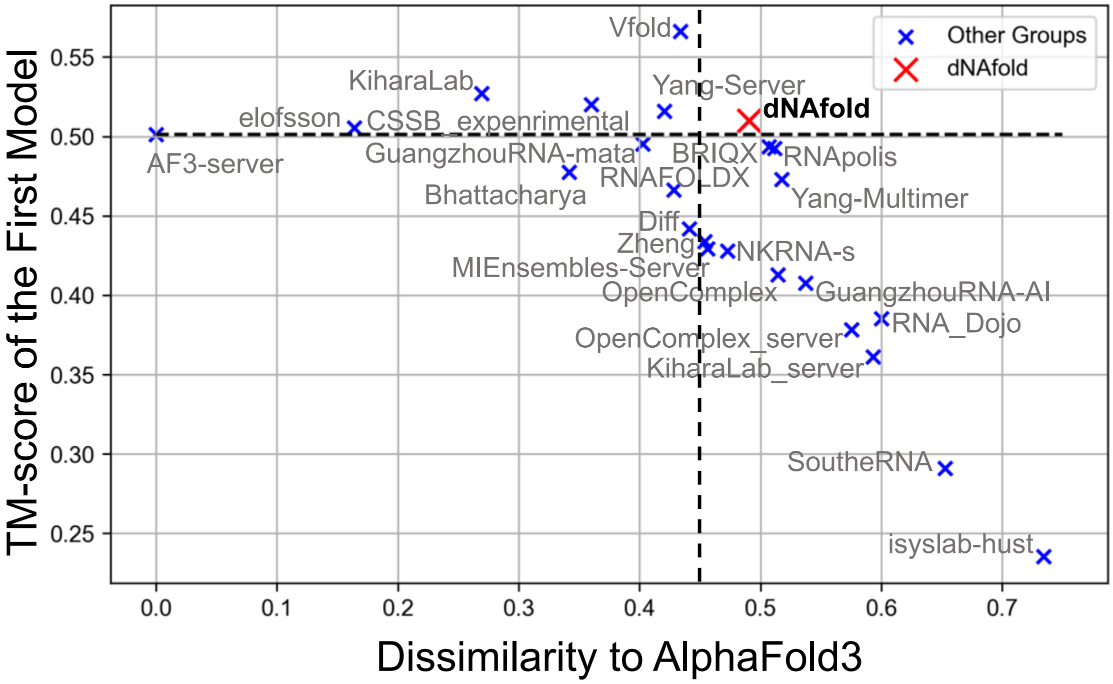


**Figure S2**. **Average TM-score of the first predicted models on 20 RNA monomer targets (< 400 Nts) by CASP16 groups versus the dissimilarity to AlphaFold3**. For each group, $dissimilarity = 1 - TM_M$ where $TM_M$ is the maximum TM-score between the first model and the five models from AlphaFold3.

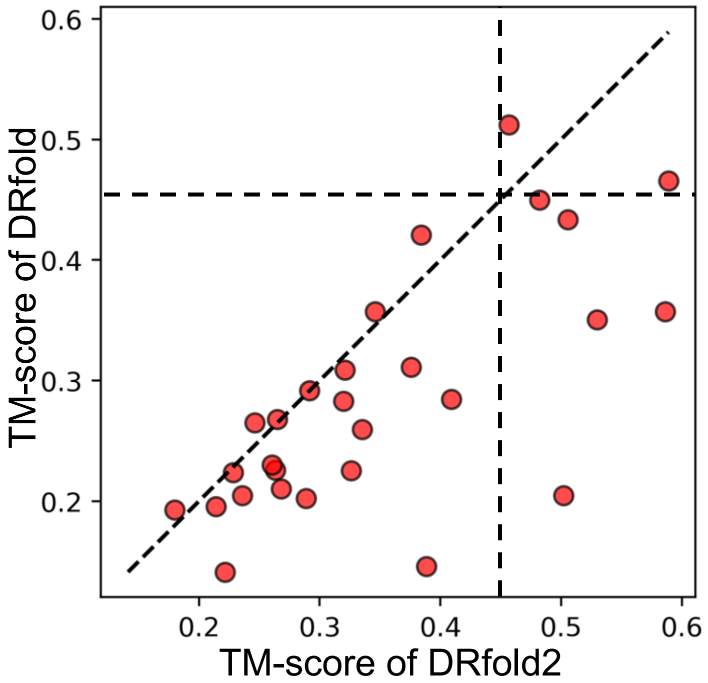


**Figure S3. TM-score comparison between DRfold and DRfold2 on the 28 test RNA structures.**
